# Bayesian phylodynamic inference of multi-type population trajectories using genomic data

**DOI:** 10.1101/2024.11.26.625381

**Authors:** Timothy G. Vaughan, Tanja Stadler

**Affiliations:** Department of Biosystems Science and Engineering, ETH Zurich, Klingelbergstrasse 48, 4056, Basel, Switzerland; Swiss Institute of Bioinformatics, Quartier Sorge - Bâtiment Amphipôle, 1015, Lausanne, Switzerland

**Keywords:** phylodynamics, particle filter, epidemiology, Bayesian phylogenetics

## Abstract

Phylodynamic methods provide a coherent framework for the inference of population parameters directly from genetic data. They are an important tool for understanding both the spread of epidemics as well as long-term macroevolutionary trends in speciation and extinction. In particular, phylodynamic methods based on multi-type birth-death models have been used to infer the evolution of discrete traits, the movement of individuals or pathogens between geographic locations or host types, and the transition of infected individuals between disease stages. In these models, population heterogeneity is treated by assigning individuals to different discrete types. Typically, methods which allow inference of parameters under multi-type birth-death models integrate over the possible birth-death trajectories (i.e. the type-specific population size functions) to reduce the computational demands of the inference. As a result, it has not been possible to use these methods to directly infer the dynamics of trait-specific population sizes, infected host counts or other such demographic quantities. In this paper we present a method which infers these multi-type trajectories with minimal additional computational cost beyond that of existing methods. We demonstrate the practicality of our approach by applying it to a previously-published set of MERS-CoV genomes, inferring the numbers of human and camel cases through time, together with the number and timing of spillovers from the camel reservoir. This application highlights the multi-type population trajectory’s ability to elucidate properties of the population which are not directly ancestral to its sampled members.

## Introduction

The strong connections between genetic diversity and population dynamics play an integral role in our ability to use modern sequencing technology to draw conclusions about populations at every scale. The field of phylodynamics offers a growing number of model-based approaches to inferring population dynamics from sequence data by way of phylogenies. While the term “phylodynamics” was originally conceived in the context of molecular epidemiology (Grenfell et al. 2004), the techniques which are now described as phylodynamic methods are applied in a diverse range of contexts from macroevolution (Gavryushkina et al. 2017) to developmental biology (Stadler et al. 2021).

One widely-used family of phylodynamic models are the so-called birth-death models (Kendall 1948) under which the population dynamics are the result of a stochastic process parameterized by individual birth and death rates. Combined with an explicit sampling process (Stadler 2010) and time-dependent rate shifts (Stadler et al. 2012), these models enable genomic inference of the past dynamics of critical epidemiological and macroevolutionary parameters, such as transmission rates, speciation rates, and migration rates.

Beyond these parameters, the dynamics of the populations themselves are potentially of interest. In epidemics these correspond to incidence and prevalence dynamics, while for macroevolutionary applications of birth-death models they correspond to species richness trajectories. Such trajectories, which summarize the dynamics of all members of a population— including those which do not belong to the phylogenetic tree relating sampled individuals—are an implicit component of all birth-death models which, at their most basic level, provide a generative description for the dynamics of a whole population as a function of rate parameters. While earlier birth-death phylodynamic inference approaches (Stadler et al. 2012; Heath et al. 2014; Gavryushkina et al. 2014) implicitly averaged over these trajectories for reasons of computational efficiency or because these trajectories were not of explicit interest, recent methods allow population trajectories to be jointly inferred with the birth-death parameters. For instance, the approach of Vaughan et al. (2019) employs a particle marginal Metropolis-Hastings algorithm to jointly infer parameters and population trajectories. This approach can be applied to nonlinear birth-death models (e.g. epidemiological models where pathogen transmission rates depend on the availability of susceptible individuals), but can be extremely demanding computationally due to its reliance on simulation-based particle filtering for likelihood evaluation at every step of the Markov chain Monte Carlo (MCMC) inference algorithm. In contrast, the more recent approach of Manceau et al. (2021) is specific to linear birth-death models where individual birth and death rates lack density dependence, but allows for more tractable inference of population history via numerical integration of sets of differential equations.

The above models and methods focus on neutrally-evolving, unstructured populations, where birth, death and sampling rates are the same for all co-existing individuals; assumptions which are inappropriate for many applications. This has spurred the development of a sub-family of birth-death phylodynamic models which allow population members to be associated with a discrete type whose two (Maddison et al. 2007) or more (FitzJohn 2012; Kühnert et al. 2016) values may influence the birth, death and sampling rates to which they are subject. These models have been applied extensively to the study of trait-dependent speciation and extinction (eg. Maddison et al. 2007; FitzJohn 2012; Rabosky 2014; Barido-Sottani et al. 2020) and to epidemiological questions, where the discrete types have been used to model the transmission dynamics of pathogens across locations (eg. Nadeau et al. 2021), host types (eg. Grear et al. 2017; Guinat et al. 2022), and pathogen strains (eg. Rasmussen and Stadler 2019; Loiseau et al. 2023).

While the applications are diverse, for existing multi-type inference methods (eg. FitzJohn 2012; Kühnert et al. 2016) only the parameters defining the type-dependent rates of birth, death and sampling of individuals, together with the phylogenetic tree relating samples can be readily inferred. This is despite the fact that—like their single-type counterparts—the multi-type birth-death models which underpin such methods describe the generation of full multi-type population dynamics, not only the phylogenetic tree relating the sampled sequences. As in the single-type case, multi-type birth-death phylodynamics inference methods implicitly average over this richer picture of the population dynamics.

Here we present an approach to performing joint Bayesian inference of the phylogenetic tree, multi-type birth-death model parameters, and ancestral lineage types, together with type-specific population trajectories. Importantly, our approach allows this inference to be done with only a relatively small increase in the computational run time of the inference relative to that of the existing multi-type phylodynamic approaches of Kühnert et al. (2016) and Scire et al. (2022). This is accomplished by “mapping” trajectories onto combinations of trees and parameters already sampled using existing MCMC techniques. The approach and its implementation are directly applicable to all models considered by Kühnert et al. (2016) and Scire et al. (2022), namely linear multi-type birth-death models with a defined but otherwise arbitrary number of types, where type-specific birth, death, migration and sampling parameters can change through time in a piecewise-constant fashion. After introducing the approach and demonstrating the correctness of its implementation, we apply the method to a published set of MERS-CoV genomes curated by Dudas et al. (2018) belonging to the 2012–2015 MERS outbreak in the Arabian Peninsula. These authors used the results of a coalescent-based ancestral-state inference method (Vaughan et al. 2014) to deduce the presence of camel to human spillover events in those lineages ancestral to the observed pathogen genomes. Our results demonstrate how the new approach enables Bayesian phylodynamic inference of host-specific prevalence dynamics, which yields information about the number and timing of camel to human spillover events in the broader population from which the samples were collected.

### New Approaches

Our approach to inferring multi-type trajectories from sequence data is based on the combination of a multi-type extension to our particle filtering approach (Vaughan et al. 2019) with a stochastic mapping approach to the placement of ancestral type changes on a tree. In this section, we describe the model assumptions, outline the new inference approach, and derive the necessary extensions to the particle filter. (While mathematical notation in this section is explained as it is introduced, table 1 provides a quick reference for the most important notation employed.)

**Table 1.**
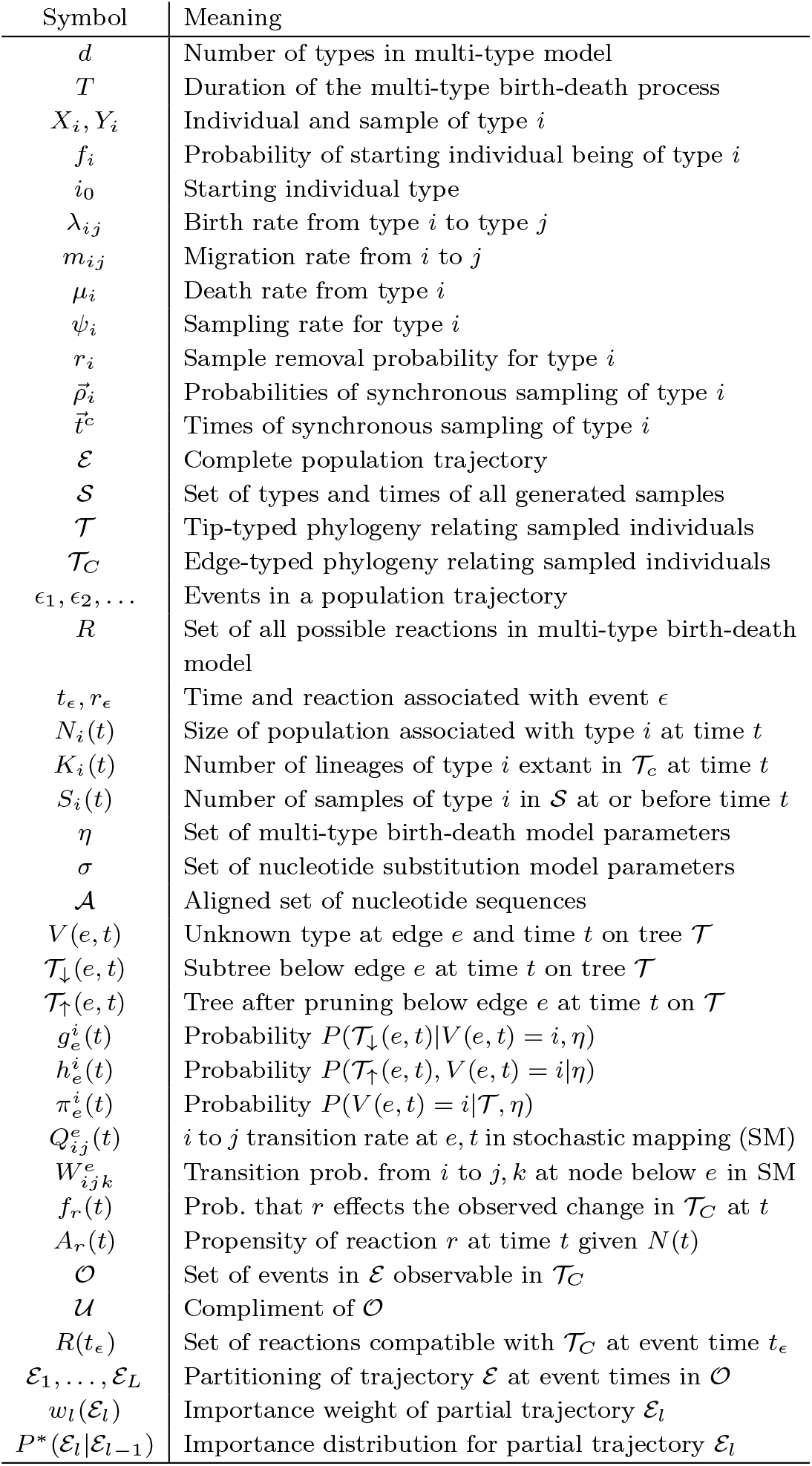
Meanings of mathematical symbols used in this paper.

#### Linear birth-death phylodynamic models

We focus here on a population composed of a discrete set of individuals, each belonging to one of a fixed number d of distinct types. This multi-type system is assumed to evolve according to a stochastic linear birth-death branching process parameterized by rates which in general vary through time in a piecewise-constant way. The process begins at time 0 with a single individual, whose type *i*_0_ ∈ [1, *d*] is chosen randomly with probability 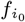. The process continues for a duration *T*. While variations of this process have been described by numerous authors over the years, we here consider the specific form described by Kühnert et al. (2016) and by Scire et al. (2022).

We introduce the notation *X*_*i*_ to represent an individual of type *i*. Additionally, we use *Y*_*i*_ to represent a sample of type *i*. The individuals comprising the population are subject to the following named reactions:

- Birth_*ij*_: An individual of type *i* gives birth to a new individual of type *j* with rate *λ*_*ij*_:

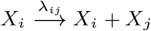
- Death_*i*_: An individual of type *i* dies at rate *µ*_*i*_:

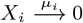
- Migrationij : An individual of type *i* changes type to *j* at rate

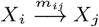
- SampR_*ij*_ : An individual of type *i* is sampled and removed at rate *ψ*_*i*_*r*_*i*_, where *ψ*_*i*_ represents the overall sampling rate and *r*_*i*_ is the probability of removal on sampling:

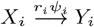
- SampNR_*ij*_ : An individual of type *i* is sampled without removal at rate *ψ*_*i*_(1 − *r*_*i*_):

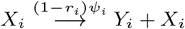

The linearity of the process is reflected in the dependence of each of these reactions on only a single reagent. The rates associated with these reactions indicate the probability per unit time of individual *X*_*i*_ undergoing the corresponding reaction at time *t*. Each of the rate parameters *λ*_*ij*_, *µ*_*i*_, … are assumed to be piecewise-constant functions of time, meaning that the process is in general time-inhomogeneous. (For clarity of exposition we do not explicitly indicate this time dependence in what follows; rather the derivatives and integrals where these functions appear should all be understood to be defined in a piecewise fashion.)

In addition to these continuous reactions, the population is subject to synchronous sampling at the times 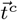. At a given synchronous sampling time 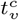, every extant lineage of type *i* is sampled with probability 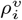, and each sampled lineage is removed from the population with probability 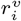. The resulting reactions are given the parameterized name SyncSample 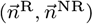, where 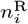 and 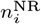 are respectively the number of removed and non-removed samples of type *i* produced by the reaction.

A complete realization of the multi-type birth-death process would be a full, binary tree recording the ancestral relationships between all events and individuals that appeared in the system. However, we are only interested those properties of a given realization about which we can reasonably learn using genetic data associated with a subsample of the population. We thus consider only these elements: the population trajectory ℰ, the times and types of all samples 𝒮, the phylogenetic tree 𝒯 relating these typed and timed samples, and the phylogenetic tree whose edges are annotated with ancestral types, 𝒯_*C*_. (In the parlance of Kühnert et al. (2016), we refer to 𝒯 as a *tip-typed tree* and 𝒯_*C*_ as an *edge-typed tree*.) In other words, we ignore the topology of those parts of the full tree not directly ancestral to the samples 𝒮.

The population trajectory ℰ is composed of the starting population type *i*_0_ together with a sequence of events *ϵ*_1_, *ϵ*_2_, …. An event *ϵ* consists of a time, t_*ϵ*_, and the corresponding reaction, r_**ϵ**_. (Synchronous sampling reactions always produce an event in the event sequence, even when no samples are generated.)

For what follows it will be helpful to define several vector-valued piecewise-constant auxiliary functions of time which can be uniquely derived from *ℰ* and 𝒯_*C*_. First among these is 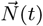, whose elements *N*_*i*_(t) represent the size of the population associated with type *i* at time *t*. Secondly we define 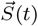, whose *i*^th^ element represents the cumulative number of samples collected from sub-population *i* up until time *t*. Finally, we define the function 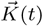, whose elements represent the number of extant lineages of 𝒯_*C*_ associated with each type at time *t*. These auxiliary functions, together with corresponding tip-typed 𝒯 and edge-typed 𝒯_*C*_ trees, are illustrated in Figure 1.

**Fig. 1:**
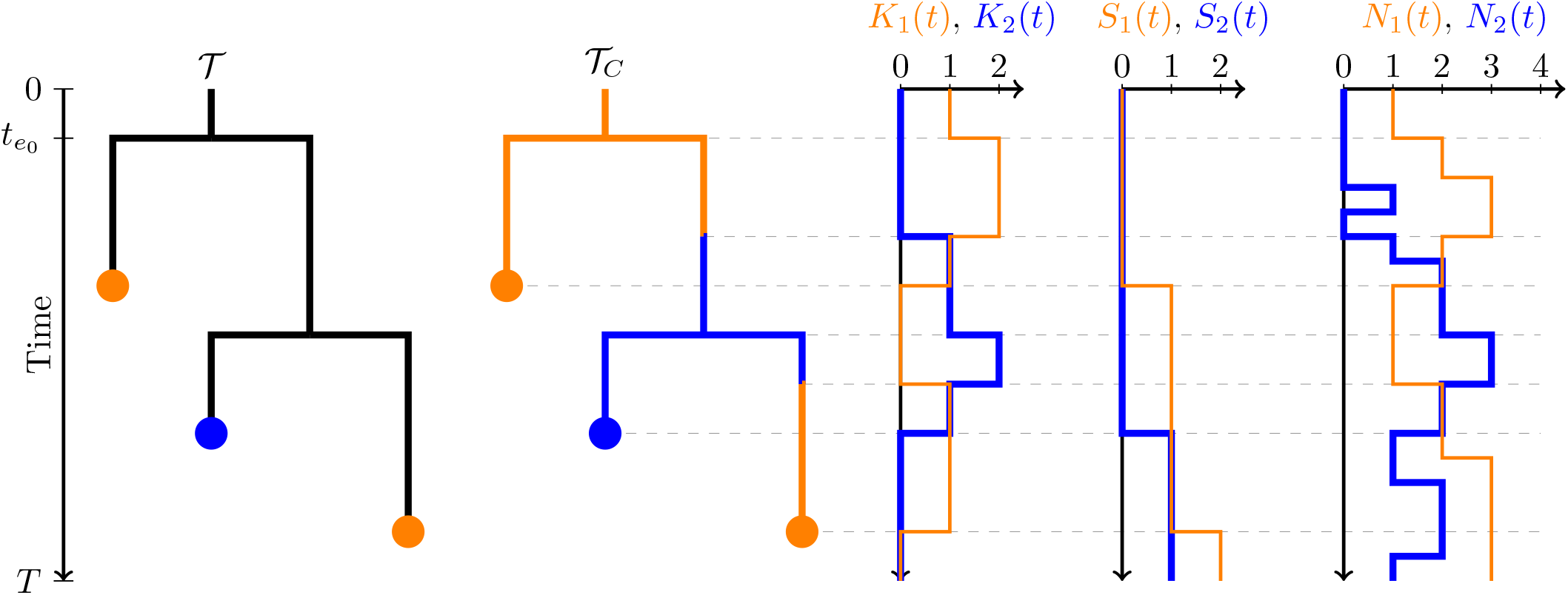
Relationship between the tip-typed tree 𝒯, the edge-typed tree 𝒯_*C*_, the multi-type lineage-through-time function 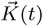, the cumulative sample count function 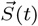 and the multi-type population size function 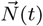, for a single realization of a multi-type birth-death-sampling model starting at time 0 and ending at time *T*. The time of the root of the tree is 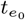.

#### Bayesian inference strategy

Our focus here is on jointly inferring population dynamics ℰ and the edge-typed tree 𝒯_*C*_ from the number, timing and types of samples *𝒮* produced by the population together with a genetic sequence alignment 𝒜. The sequences are assumed to have evolved down the reconstructed tree 𝒯 according to a standard Markovian nucleotide substitution model. That is, we seek to characterize the distribution P (ℰ, 𝒯_*C*_, *σ, η*| *𝒜, 𝒮*), where *σ* and *η* are the parameters of the nucleotide substitution and birth-death models, respectively.

One approach to this problem would be to use Markov chain Monte Carlo (MCMC) directly to jointly sample the edge-typed tree 𝒯_*C*_, the multi-type population trajectory *E*, and all model parameters. However, this approach would likely prove impractical due to the extremely large dimensionality of the state space. (While MCMC algorithms for directly sampling edge-typed trees under a variety of models are in use (eg. Beerli and Felsenstein 2001; Ewing et al. 2004; Vaughan et al. 2014; Kühnert et al. 2016), they are known to suffer from convergence issues which motivate the use of alternative schemes (Hey and Nielsen 2007; De Maio et al. 2015; Müller et al. 2018). The addition of the highly-dimensional trajectory variable ℰ to this space would likely make this problem worse.)

Instead, the approach that we advocate here is analogous to the general approach championed by Hey and Nielsen (2007) to improve the efficiency of inference of migration rates under the isolation-migration coalescent model, and applied more recently to the inference of ancestral types under the multi-type birth-death model by Freyman and Höhna (2018). This approach still involves MCMC, but to sample a marginalized posterior of lower dimensionality that includes neither ancestral types nor population trajectories. Instead, these latent variables are independently drawn from their conditional probability distributions given the MCMC-sampled tip-typed trees and birth-death parameters.

The following factorization of the joint posterior is central to our approach:

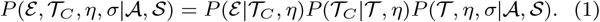

The approach involves three steps, each corresponding to a term in the factorization:

1. sampling 𝒯, *η, σ* ~ *P* (𝒯, *η, σ*| A, *𝒮*) using the MCMC algorithm described and used by Kühnert et al. (2016) and Scire et al. (2022),
2. sampling 𝒯_*C*_ ~ *P* (𝒯_*C*_ | 𝒯, *η*) using a stochastic mapping algorithm for ancestral state reconstruction similar to the approach developed by Freyman and Höhna (2018), and
3. sampling ℰ ~ *P* (ℰ | 𝒯_*C*_, *η*) using a particle filtering algorithm. These three steps allow us to produce trajectories, phylogenies and parameters sampled from the joint distribution given by Eq. 1.

On their own, the trajectories are distributed according to their marginal posterior *P* (ℰ | 𝒜, 𝒮).

A key advantage of this approach is that the second and third steps need only be applied to a small subset of the samples of 𝒯, *η, σ* produced by the MCMC chain. This is because the sequence of states produced by MCMC algorithms are usually highly autocorrelated, meaning that a small subset of the states visited can characterize the target distribution as well as the full chain. While there is no well-defined consensus on the number of “effectively independent” samples required to produce useful characterizations of phylodynamic posteriors (although for parameter estimates it is common to treat values above 200 as acceptable), it is clear that this number is usually many orders of magnitude smaller than the number of iterations needed by the MCMC algorithm to achieve it. Thus, the ability to apply steps 2 and 3 to this subset means that these components of the approach are almost cost-free relative to the far larger computational cost of the MCMC portion of the algorithm.

As stated above, the MCMC algorithm we use for step 1 above is precisely that presented by Kühnert et al. (2016), thus we do not describe it further here. The following sections describe the algorithms used to sample from the conditional distributions required for steps 2 and 3.

#### Stochastic mapping of ancestral types

Here we describe the approach we take to generating samples from P (𝒯_*C*_ | 𝒯, *η*) which is based on the same principle as the approach described by Freyman and Höhna (2018), namely a post-order (leaves to root) numerical integration followed by a pre-order (root to leaves) lineage type simulation. We present here a succinct derivation of this algorithm in terms of an appropriately conditioned master equation for the type change process along tree edges. This algorithm is applicable to all trees which can be produced by the multi-type birth-death-sampling model of Kühnert et al. (2016) and Scire et al. (2022)—including non-ultrametric trees and trees with sampled ancestor nodes (Gavryushkina et al. 2014).

Consider a particular time *t* on a particular edge *e* of the tree 𝒯. We use this point (*e*, *t*) to divide the tree into two components: 𝒯_*↓*_(*e*, *t*) being the clade descending from this point, and 𝒯_↑_(*e*, *t*) being the portion of the tree remaining when this clade is removed.

By definition, these components are perfectly complementary in the sense that 𝒯 = 𝒯_*↓*_(*t*, *e*) ∪ 𝒯_↑_(*e*, *t*).

Defining *V*(*e*, *t*) to be the (unknown) ancestral type value at (*e*, *t*), we can express the marginal probability that this trait takes a particular value *i* as

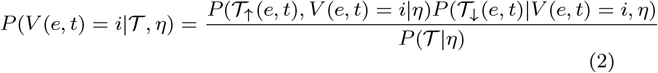

Noting that 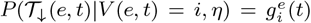 where 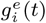 is the _*i*_ partial tree probability function considered by Kühnert et al. (2016), and using the notation 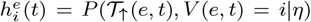 and 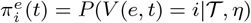, we can write this as

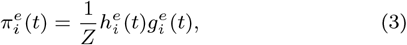

where *Z* = *P*(𝒯 |*η*) is the normalization constant.

Our goal is to determine a stochastic process which produces realizations of *V*(*e*, *t*) for every point in the tree. To do this, we use the above expansion to derive the master equation for the marginal ancestral type distribution by directly differentiating this equation:

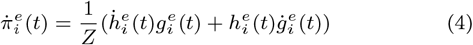

As described by Kühnert et al. (2016) and Scire et al. (2022), the time derivative for the partial tree probability 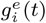 along an edge is given by the following system of ordinary differential equations (ODEs):

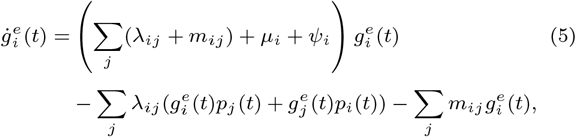

where *p*_*i*_(*t*) is the probability that an individual of type *i* alive at time *t* yields no sampled descendants, which follows its own ODE also described by Kühnert et al. (2016) and Scire et al. (2022). (The synchronous sampling probabilities appear only in the boundary conditions of these ODEs, as they do not affect the continuous-time evolution of the tree probabilities.)

We now make use of the normalization constraint that 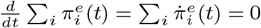 to give

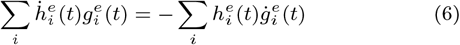

Substituting equation 5 into the right-hand side and performing a change of variables on the summation index, this becomes

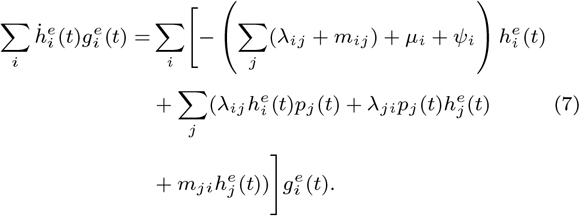

For this equation to hold for arbitrary 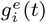, the summation kernels must be equal. Thus

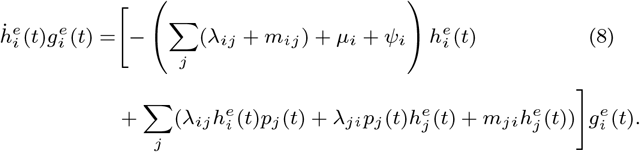

Substituting this, together with equation 5 back into equation 4 yields

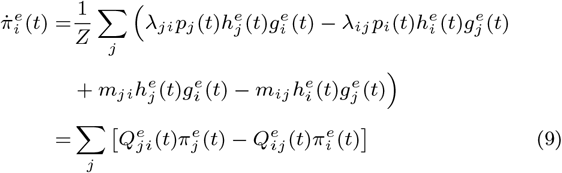

where we have defined

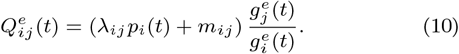

We thus find that, conditional on the tree and parameter values, the ancestral type evolves down the tree according to a continuous-time Markov process with time-dependent transition rates 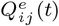. (The presence of piecewise-constant variation in the multi-type birth-death rate functions simply induces corresponding piecewise-constant variation in these transition rates.)

In addition to changes along edges, state changes can occur at internal nodes when the off-diagonal elements of the rate matrix *λ*_*ijk*_ are non-zero. Using simple arguments (see Supplemental Text) we can identify the transition probabilities for state changes at an internal node below edge e given a parental type i, 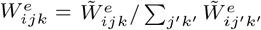, where

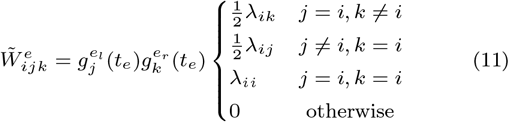

Here *t*_*e*_ represents the time of the node below edge *e*, while *e*_*l*_ and *e*_*r*_ represent the left and right child edges of the node.

Since we can pre-compute the components of these rates and transition probabilities by numerically solving the differential equations given by Kühnert et al. (2016), we can directly sample the state changes by (a) sampling the starting state from its marginal posterior, then (b) simulating this Markov process forwards in time down the tree.

Sampling the appropriately conditioned starting state can be done directly by noting that at the start of the root edge 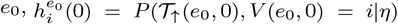 reduces to *P* (*V*(*e*_0_, 0) = *i*|*η*) = *f*_*i*_, and thus that we can compute the marginal probabilities for each possible starting state 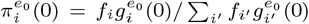 directly.

Simulating the process and hence drawing a sample from *P*(𝒯_*C*_ | 𝒯, **η**) can be done using any algorithm capable of simulating trajectories from a continuous-time Markov chain with time-dependent rates. The approach we use, given as algorithm 1, is a simple extension of the original Stochastic Simulation Algorithm (Gillespie 1976).

##### Algorithm 1

Stochastic mapping algorithm. The notation *t*_*e*_ represents the time of the youngest node attached to edge *e*, ChildCount(*e*) represents the number of its child edges, Child(*e*) is the single child edge when ChildCount(*e*)=1, and LeftChild(*e*) and RightChild(*e*) are the left and right child edges when ChildCount(*e*)=2.

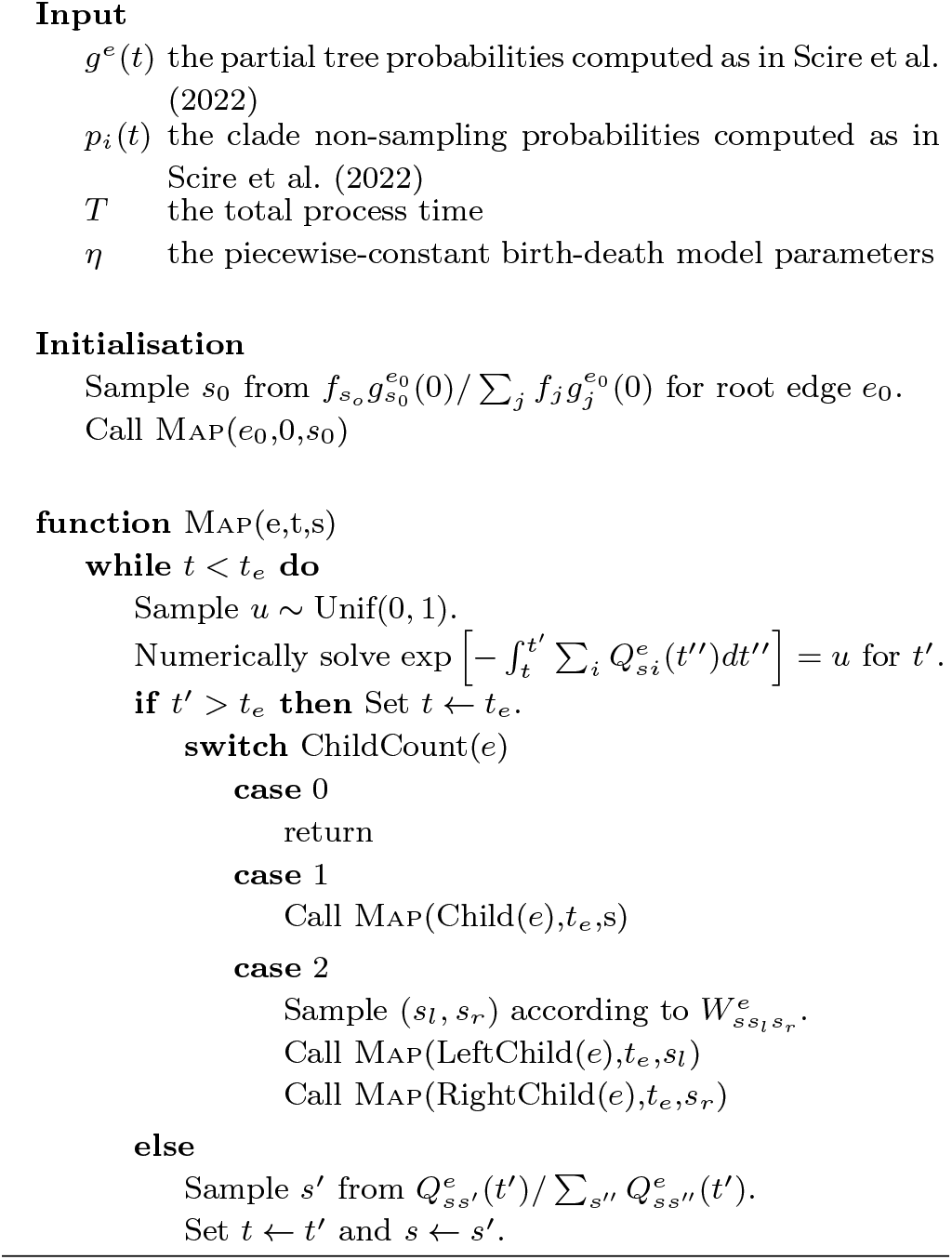

#### Importance sampling trajectories

While it is straightforward to produce samples of the multi-type trajectories from the unconditioned distribution *P*(*ℰ* |**η**) through direct stochastic simulation (Gillespie 1976), our goal is to sample ℰ from the conditional distribution *P* (ℰ | 𝒯_*C*_, **η**). We will bridge this gap using a particle filtering approach that we will present in the next section. Before describing that approach however, we will firstly consider the more fundamental problem of developing an importance sampler for the conditional distribution, with the goal of providing an intuitive basis for understanding the particle filtering algorithm presented in the next section.

A naïve importance sampler might use direct simulation to sample an ensemble of *M* ≫ 1 trajectories directly from *P*(*E*|*η*) and weight each trajectory using the edge-typed tree probability *P* (𝒯_*C*_ | *ℰ*). However, literal application of this method is not practical, since *P* (𝒯_*C*_ | ℰ ^(*m*)^) will be zero for any ℰ ^(*m*)^ ~ P (ℰ |**η**). This is because the tree density is only non-zero when the real-valued times of all events represented on the tree coincide precisely with the times of events in ℰ ^(*m*)^.

For this reason, we instead simulate trajectories from the modified distribution *P* ^∗^(ℰ|**η**). This distribution corresponds to a process similar to *P* (ℰ|**η**) but with the important distinction that all events contained in the edge-typed tree 𝒯_*C*_ occur with probability 1, and any events that are incompatible with 𝒯_*C*_ are forbidden. Weighting these trajectories with the unnormalized weight function

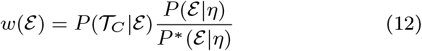

produces our desired target distribution, which can be seen by noting that:

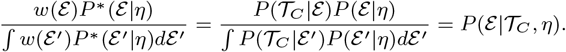

To compute these weights, we need to specify three distributions: the trajectory probability *P* (ℰ|**η**), the importance trajectory distribution *P* ^∗^( ℰ|**η**) and the tree probability *P* (𝒯_*C*_ |*E*).

The trajectory distribution follows directly from the definition of the birth-death process, and can be written

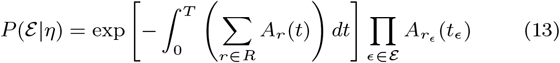

where R represents the set of all possible continuous reactions in the model (i.e. excluding synchronous sampling reactions). For such reactions, *A*_*r*_(*t*) is the piecewise-constant propensity of reaction *r* at time *t* (e.g. for 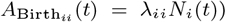, with the temporal dependence stemming from both population size variation and any piecewise-constant variation in rate functions. For synchronous sampling reactions on the other hand, 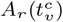 is the multinomial probability of the particular removed/non-removed synchronous sampling outcome at the sampling time 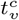, that is:

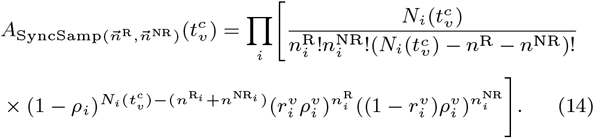

In both cases, *r*_*ϵ*_ is the reaction involved in event *ϵ*, and *t*_*ϵ*_ is the time of that event.

The probability of the edge-typed tree can be similarly decomposed:

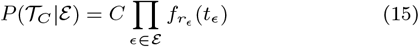

where *C* is a normalizing constant and

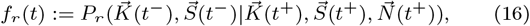

the probability for a reaction *r* to effect a specific instantaneous change in the lineages-through-time function and cumulative sample count function at time *t*. The times *t*^−^ and *t*^+^ refer to the times immediately before and after *t*. The functions 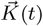 and 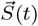 are the lineages-through-time and cumulative sample count functions for 𝒯_*C*_, and 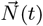 is the population size function for ℰ —as illustrated in Figure 1.

The normalizing constant C accounts for the fact that, in general, many unique tree topologies correspond to a given pair of lineages-through-time and cumulative sample count functions. Since the number of compatible topologies affects only the overall normalization of the weight function, it can be safely ignored.

The probabilities *f*_*r*_(*t*) are derived by considering the number of ways that lineages belonging to the tree can be involved in a given reaction, and are given below for each of the reactions in our model. (In the following, 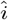 and 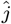 represent unit vectors of dimension d where the *i*^th^ or, respectively, *j*^th^ element is 1 and all other elements are 0.)

- Single-type birth reactions:

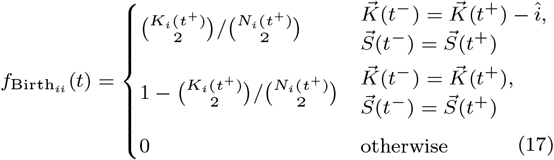
- Between-type birth reactions:

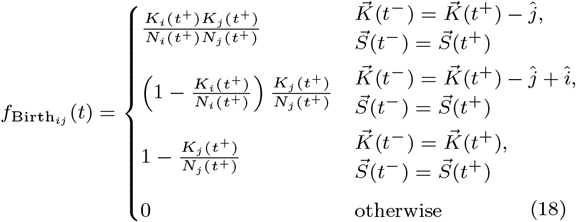
- Migration reactions:

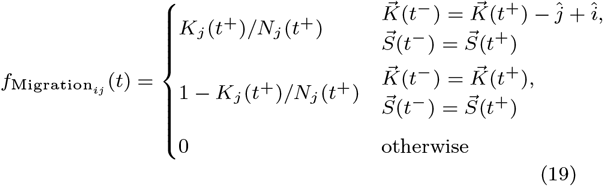
- Death reactions:

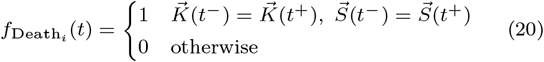
- Sampling reactions with removal:

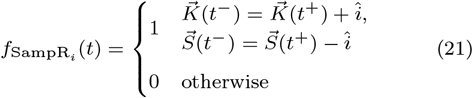
- Sampling reactions without removal:

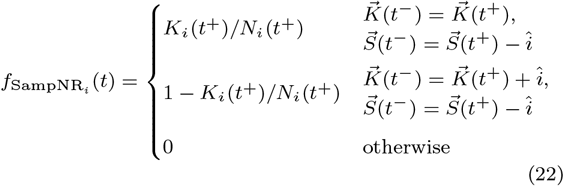
- The probability due to synchronous sampling events is a product of hypergeometric distributions for the number of sampled ancestors of each type produced by a synchronous sampling reaction at a given time:

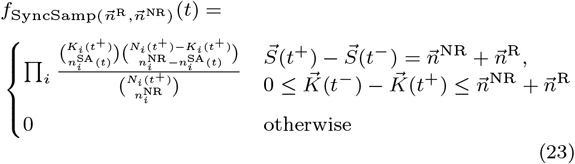

(Here 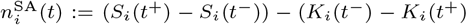), i.e. the difference between the number of type-i samples generated at time t and the number of new type-*i* lineages appearing at the same time, represents the number of “sampled ancestors” of type *i* generated by the 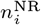 unremoved samples.)

Note that these tree event probabilities are a superset of those used by the particle filtering algorithm presented by Vaughan et al. (2019).

In order to express the importance distribution, we partition the trajectory ℰ = 𝒰 ∪ 𝒪, where 𝒪 includes only the events occurring at exactly the same time as events on the tree (“observed” events), and 𝒰 contains the remaining (“unobserved”) events. The importance distribution corresponds to a modified birth-death process in which unobserved events occur at rate *A*_*r*_(*t*)*f*_*r*_(*t*), while observed events occur with probability 1 and with a reaction *r*_**ϵ**_ chosen with probability 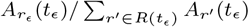 from the set *R*(*t*_**ϵ**_) of reactions (including synchronous sampling reactions) compatible with the tree at the observed event time *t*_**ϵ**_. This modified process ensures that (a) only those unobserved events compatible with the tree are included, and that (b) observed events occur with probability 1 and are associated with reactions also compatible with the tree. The probability for a trajectory generated by this process is written:

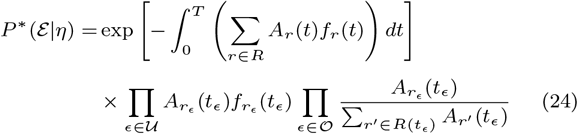

Substituting this, together with the expressions for the tree and trajectory probabilities, into equation (12) yields the importance weight function:

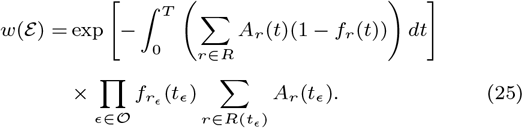

One could thus sample trajectories from *P* (ℰ| 𝒯_*C*_, **η**) by drawing a large number of trajectories directly from *P* ^∗^( ℰ|*η*), computing their associated weights using *w*(ℰ), then drawing from this weighted distribution. However, since the weights are assigned only once the trajectories have been completely simulated, the weighted distribution will likely include many trajectories with relatively low weights. This will yield weighted ensembles a low effective sample size However, while this importance sampling strategy will produce the correct distribution in the limit of an infinitely large trajectory ensemble, it will likely perform poorly in practice where we are restricted to finite ensembles of trajectories. The reason for this is that trajectories produced by this importance sampler above are only assigned weights once they have been completely simulated, potentially leading to very low effective sample sizes for the weighted ensemble.

#### Particle filtering algorithm for multi-type trajectories

Given these considerations, we therefore employ a widely-used strategy for improving the effective sample size of the weighted ensemble known as particle filtering. (Refer to Cappe et al. (2007) for a good overview of this approach.) This involves subdividing the single importance sampler above into a number of intermediate importance samplers, with resampling steps in between. This periodic resampling helps keep disparities between trajectory weights in the ensemble manageable.

We apply this here by subdividing the trajectory ℰ into portions ℰ_1_, …, ℰ_*L*_, where each ℰ_*l*_ includes only those trajectory events with times satisfying *t*_*ϵ*_ *∈* (*t*_*l−*1_, *t*_*l*_] and where *t*_*l*_ correspond to the times of the *L* = |𝒪| events observed on the tree. We similarly partition the set of unobserved events 𝒰 into 𝒰_1_, …, 𝒰_*L*_. Since (a) the stochastic process described by the importance distribution *P* ^*\**^(ℰ|*η*) is Markovian, and (b) the weight function *w*(ℰ) has a simple product form, we can write

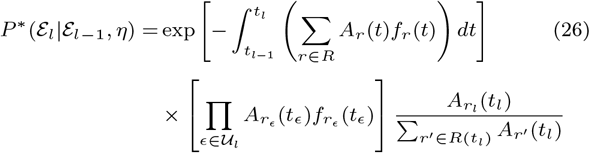

and

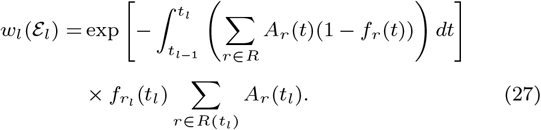

where we have used *r*_*l*_ to refer to the reaction corresponding to the observed event ending interval *l*.

Our particle filter involves initializing an ensemble of trajectories (particles) based on the initial type defined by the multi-type tree 𝒯_*C*_and then alternating between (1) propagating the ensemble to the next interval using stochastic simulation to sample from *P* ^*\**^(ℰ| ℰ_*l−*1_, *η*) defined in Eq. (26), and (2) resampling the ensemble according to the weights *w*_*l*_(ℰ_*l*_) defined in Eq. (27).

Loosely speaking, each propagation step uses importance sampling to characterize the portion of the conditioned distribution associated with the corresponding interval. By applying these smaller importance sampling steps sequentially and interspersing them with resampling steps, the particle filter keeps the variance of the particle weights from becoming unmanageable.

The full procedure is described in Alg. (2), and—in the large particle number limit—produces a single multi-type trajectory drawn from the true conditional distribution of trajectories given an edge-typed tree and corresponding multi-type birth-death parameters. This algorithm can be regarded as special case of Algorithm 3 (particle filtering) in Cappe et al. (2007). However, as noted in our previous paper (Vaughan et al. 2019) and discussed recently in more detail by King et al. (2022), when applied to such a genealogical process the weighted distribution of trajectories is only interpretable as a true conditional distribution given the tree once the entire tree has been considered.

## Results

### Implementation

The algorithms described were implemented as an extension to the Bayesian phylogenetic inference platform BEAST 2 (Bouckaert et al. 2019). Specifically, these algorithms were implemented in the package BDMM-Prime, which is a recent revision of the original BDMM package published by Kühnert et al. (2016). Beyond stochastic mapping and multi-type trajectory inference, this package also makes use of efficiency and numerical stability improvements introduced by Scire et al. (2022) and the analytical single-type functionality of the earlier BDSKY (Stadler et al. 2012) package, meaning that all analyses possible in these earlier packages are accessible within BDMM-Prime. It also includes an extensive and redesigned graphical interface, making it possible for users to configure more complex analyses compared with the earlier implementations. The new package is free software and is distributed under the GNU General Public License version 3. The source code is available from its GitHub repository (https://github.com/tgvaughan/BDMM-Prime), and a comprehensive online manual including a tutorial demonstrating its use can be found at the project website (*https://tgvaughan.github.io/BDMM-PriMe*).

#### Algorithm 2

Particle filtering algorithm used to sample a trajectory from *P* (*ℰ*| 𝒯_*C*_, *η*).

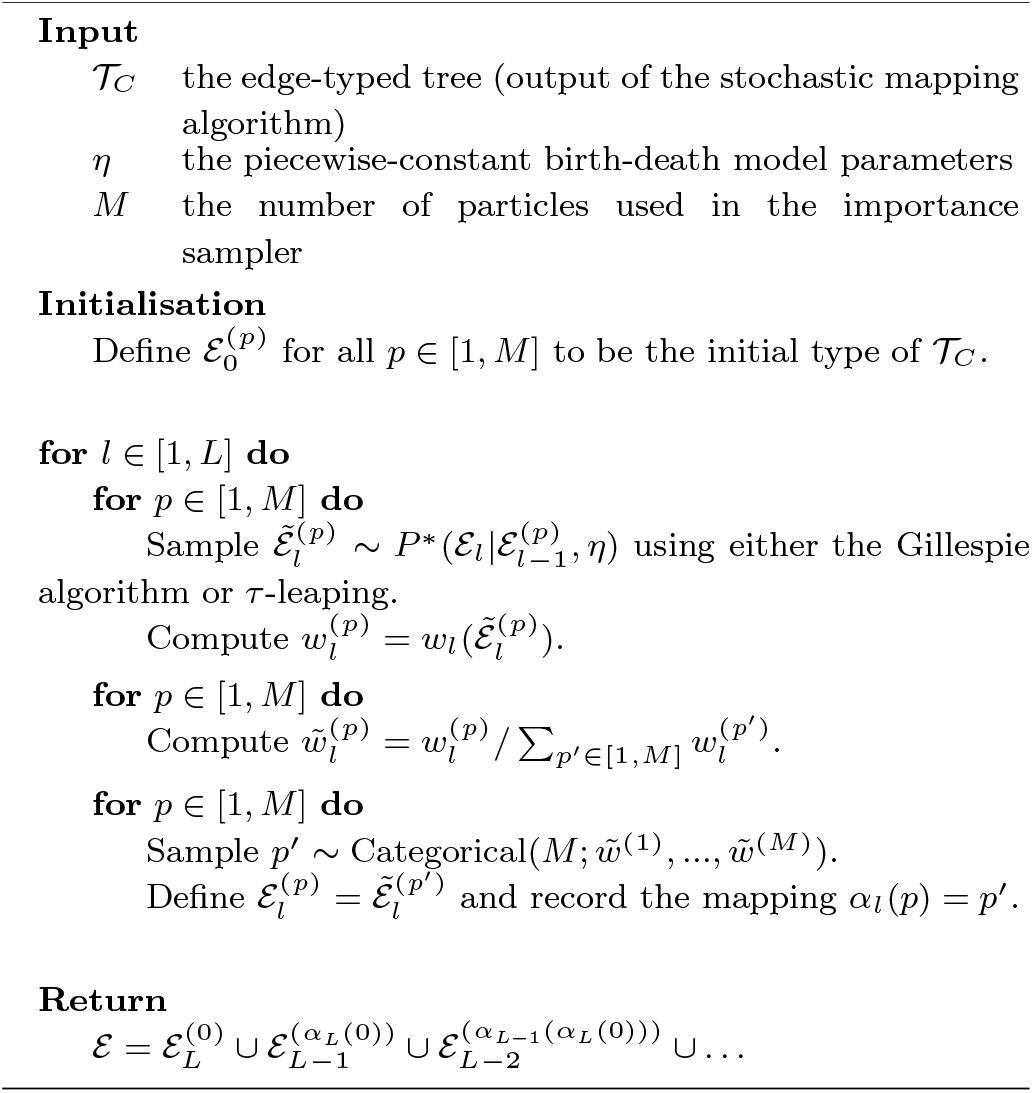

### Trait mapping validation tests

To empirically validate the implementation of the stochastic mapping approach presented as algorithm 1, we considered a family of three-type models defined by a fixed process length of 5 time units and a set of piecewise-constant rate functions defined on this domain, all with a rate shift at time 2.5 following the start of the process. From this family of models, we drew an ensemble of 10^4^ specific sets of model parameters *η* from a joint distribution *P* (*η*) composed of the following parameter-specific distributions:

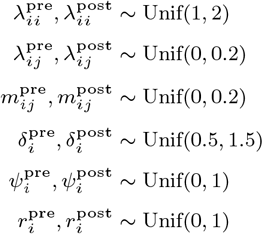

Here “pre” and “post” indicate parameter values before and after the rate shift time. We consider all combinations of *i, j ∈* 1, 2, 3 with the constraint that *j ≠ i*.

For each *η* thus sampled, we produced a corresponding edge-typed tree 𝒯_*C*_from the prior distribution *P* (𝒯_*C*_|*η*) using direct simulation. From this we produced a corresponding tip-typed tree 𝒯 by removing the ancestral type information. We then used the stochastic mapping algorithm to sample a new edge-typed tree 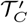.In the case that the stochastic mapping algorithm implementation correctly samples from *P* (𝒯_*C*_|𝒯, *η*), the resulting sample distributions of 𝒯_*C*_and 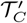 should be indistinguishable, due to the following identity:

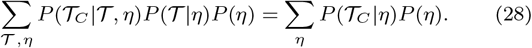

Figure 2 illustrates a comparison between two sets of summary statistics computed from these sample distributions: the total number of each of the six possible type transitions along each tree, and the total edge lengths associated with each type. Comparisons are made by considering empirical quantiles (10% through 90%) derived from each summary statistic distribution. The close agreement between the quantiles for the statistics derived from the 𝒯_*C*_ ensemble and those derived from the 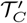 ensemble provides strong evidence for the correctness of our algorithm and its implementation.

**Fig. 2:**
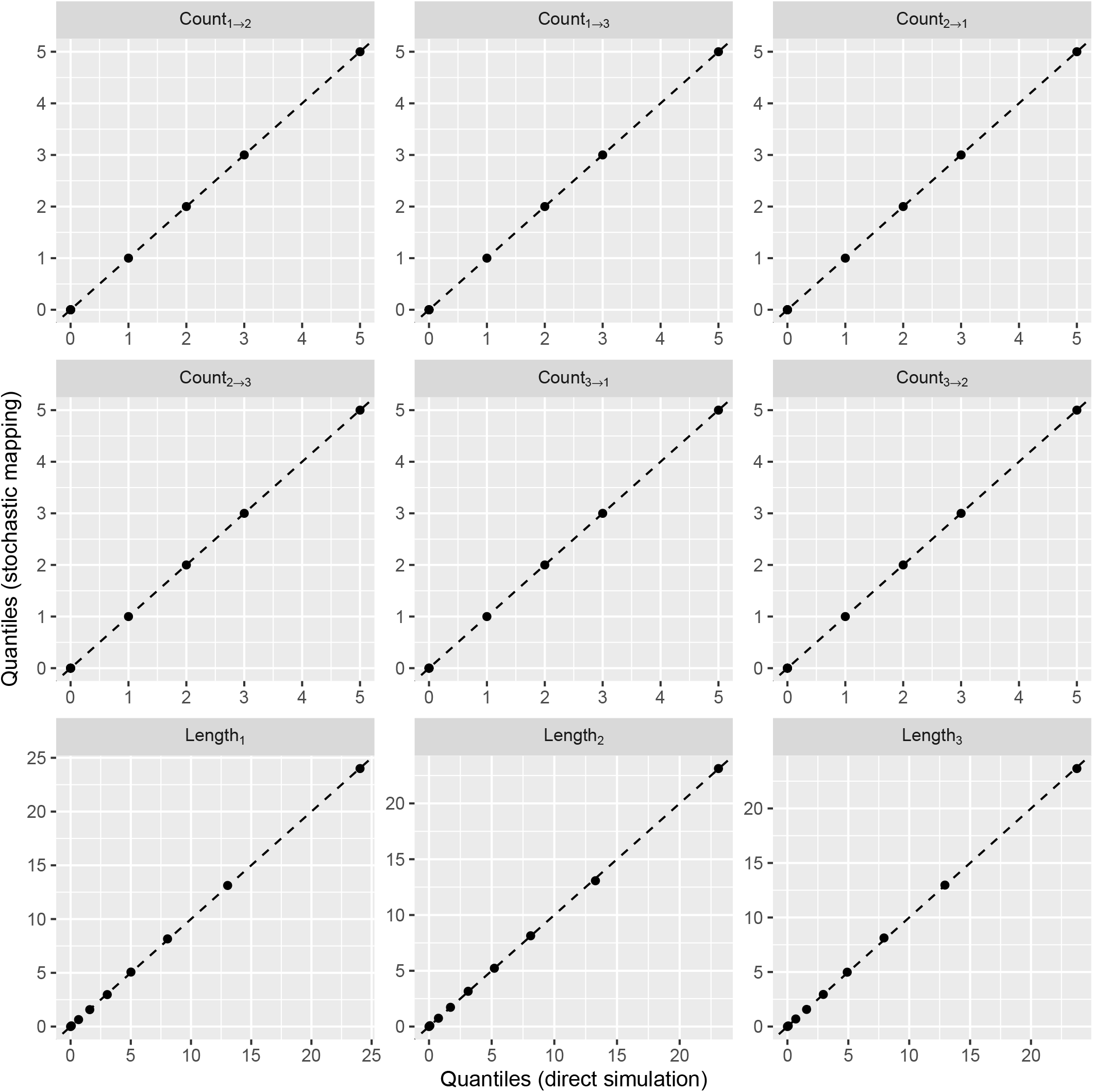
Quantile-quantile plots comparing the distributions of summary statistics for the edge-typed trees belonging to the original simulated distribution (horizontal axes) to the equivalent distributions computed from the edge-typed trees generated by the stochastic mapping algorithm (vertical axes). The statistic Count_*i→j*_ represents the number of transitions from type *i* to type *j* recorded by the edge-typed tree, while Length_*i*_ represents the total edge length associated with type *i*.

### Trajectory sampling validation tests

We took a similar approach to validating the implementation of the trajectory particle filtering algorithm. We simulated an ensemble of 10^4^ sets of parameters from the same three-type family of multi-type models defined by the same *P* (*η*) used in the stochastic mapping validation of the previous section. For each set of parameters *η*, we used direct simulation to sample a corresponding edge-typed tree 𝒯_*C*_ and trajectory ℰ. For each tree in the resulting ensemble we additionally used the particle filter with 10^3^ particles to sample ℰ′ from *P* (ℰ|𝒯_*C*_, *η*). Since

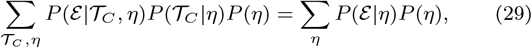

the two sample distributions for ℰ′ and ℰ should be identical, provided the particle filter used for the conditional sampling from *P* (ℰ| 𝒯_*C*_, *η*) is correct.

Figure 3 shows a comparison between these two empirical trajectory distributions. Specifically, it illustrates comparisons between the empirical quantiles (10% through 90%) of the type-specific population sizes derived from each of the two ensembles at each of 51 evenly-spaced times between 0 and 5 (i.e. the start and end of the simulation period). That none of these quantile-quantile comparisons show any meaningful difference indicates that the sample distribution derived using the particle filter indeed matches the corresponding distribution generated via direct simulation.

**Fig. 3:**
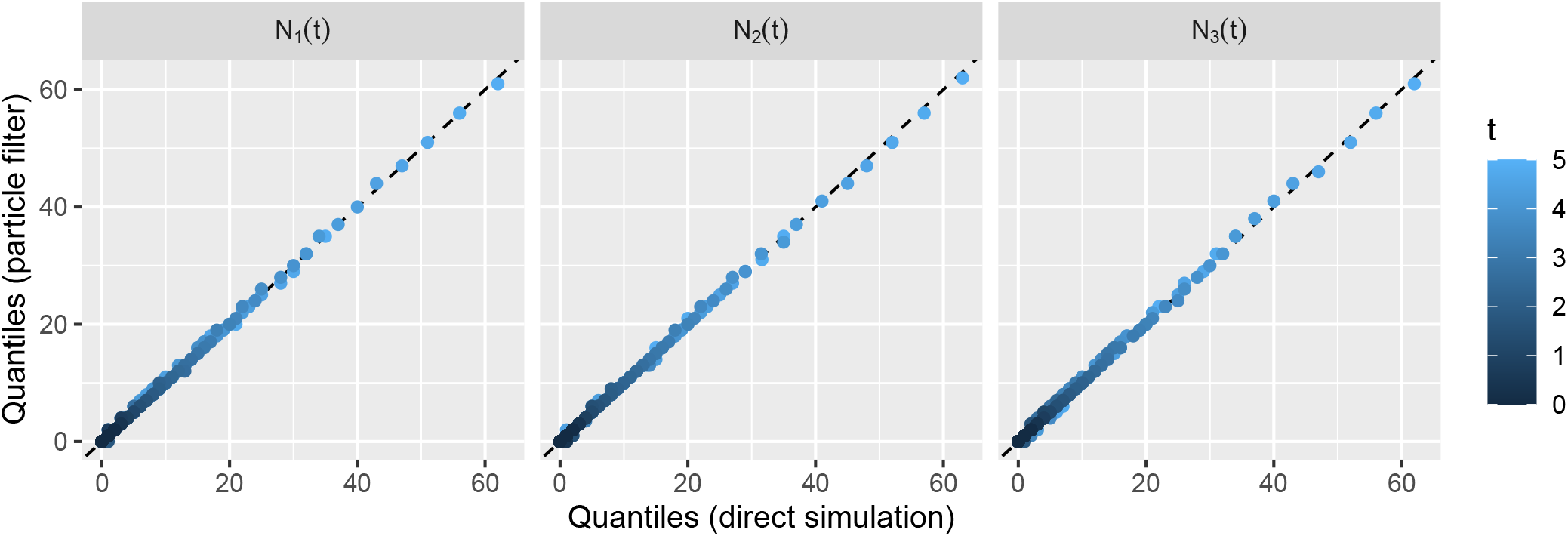
Comparisons between empirical quantiles of distributions of type-specific population sizes derived from trajectories generated by direct simulation and those generating using the particle filter, averaging over a broad family of three-type multi-type models. Each panel shows an overlaid series of quantile-quantile plots comparing population size quantiles (10% through to 90%) for a type-specific subpopulation at a 51 evenly-spaced times spanning the simulation period. The close proximity of all points to the dashed diagonal indicates equivalence between the sample distributions, which we expect for a correct implementation.

We also simulated a second ensemble of 10^4^ edge-typed trees and multi-type trajectories under a similar but distinct family of models in which the continuous sampling was replaced with a pair of contemporaneous sampling events at times 4 and 5 following the start of the process. Details of this model and the comparison between the corresponding trajectory ensembles are shown in supplemental figure S1. This also shows agreement between the distributions, further supporting our claim that the algorithm and its implementation function correctly.

### Approximating the computational burden

Rigorously assessing the time complexity of stochastic algorithms is non-trivial. However, we can readily develop an approximate understanding of the main influences on computation time of the algorithms we have presented. Firstly, the stochastic mapping algorithm relies on numerical integration of equation (5) along the tree edges, followed by stochastic simulation of the type changes using a stochastic simulation algorithm, both of which scale with the sum of all edge lengths of the tree (the total edge length). Thus we expect the calculation time for the stochastic mapping algorithm as a whole to also scale linearly with this total edge length.

The trajectory sampling algorithm, on the other hand, involves stochastic simulation of *M* candidate birth-death trajectories (i.e. particles) within each of the distinct intervals defined by the timing of features (including type changes and divergence times) on the input edge-typed tree 𝒯_*C*_. Assuming roughly constant rates, the expected number of these features scales roughly with the total edge length of the tree. We thus expect the run time of the trajectory sampling algorithm to increase linearly with both the total edge length and the number of particles used by the importance sampler.

Supplemental figures S2 and S3 respectively illustrate the calculation duration distributions of the stochastic mapping and trajectory simulation components of the validation studies reported in the previous subsections. These figures clearly show the approximately linear dependence of the time complexity of both of these calculations on the total edge length of the input trees. These figures also give an indication of the absolute time needed by these calculations in practice: for trees under 1000 leaves the stochastic mapping algorithm tends to take less than 100 ms, while the trajectory sampling algorithm requires on the order of a second when 10^3^ particles are used. Keeping in mind that the stochastic mapping and trajectory simulation algorithms are intended to be applied only to hundreds or at most thousands of tree and parameter combinations produced by multi-type birth-death MCMC analyses—which often require days to complete—such run times are practically negligible.

### Application: MERS in Camels and Humans

Following validation, we applied our approach to study the zoonotic transmission dynamics of the Middle East respiratory syndrome coronavirus (MERS-CoV). This virus was first isolated in 2012 (Zaki et al. 2012) and has been responsible for localized outbreaks of a respiratory disease, primarily in or associated with the Arabian Peninsula, with a high rate of mortality (World Health Organization 2022). While bats were initially believed to be the primary non-human reservoir, the pathogen was subsequently found to be endemic in camels in the region (see de Wit et al. (2016) for a comprehensive review).

We focused on the publicly-available MERS-CoV genomic dataset curated and used by Dudas et al. (2018) in their investigation of camel to human spillover dynamics. In that paper, the authors used a structured coalescent model (Vaughan et al. 2014) to infer the timing and number of spillover events of the pathogen from camels to humans. Our goal here was to analyze the same data using a multi-type birth-death model to jointly infer the two-host infection dynamics in the larger population, enabling direct quantification of the spillover dynamics.

The dataset is comprised of an alignment of 274 MERS-CoV genomes, including 100 genomes from camel hosts and 174 from humans. To avoid problems due to the recombination, we followed the approach taken by Dudas et al. (2018) in one of their analyses of considering only sites 21001–29364. We employed the nucleotide substitution model of Hasegawa et al. (1985) allowing for gamma-distributed site to site rate heterogeneity (Yang 1994). We assumed a relaxed clock model (Douglas et al. 2021) with an informative prior on the mean rate based on the study of MERS-CoV evolution by Zhang et al. (2016).

We performed inference under a two-type birth-death model in which the two types correspond to the two host species, and for simplicity assumed that the initial infected host was a camel. Following Kühnert et al. (2016) we used the following epidemiological parameterization of the multi-type birth-death model. Assuming lineage removal on sampling (i.e. *r*_*i*_ = 1), this parameterization can be expressed in the following way:

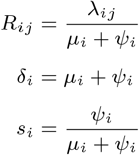

Here diagonal elements of *R*_*ij*_ represent the host-specific effective reproductive number, while off-diagonal elements generalize this quantity to describe cross-type (here zoonotic) transmission events. The parameter *δ*_*i*_ is the so-called “become uninfectious” rate, which is the net rate at which infected hosts are removed from their respective infectious compartments.

All parameters were assumed constant through time, with the single exception of the host-specific sampling proportions *s*_*i*_, which were assumed to be zero for all times outside of the interval between the first and last included sample from each host type. Prior distributions are shown in Table 2. These include informative priors which penalize extreme reproductive numbers. Additionally, they place an upper bound on the sampling proportion parameters to reflect the impossibility that our small sample of sequences represented a large proportion of the infected human or camel populations.

**Table 2.** Parameter priors used in the analysis of the MERS-CoV data. Subscripts *i* and *j* specify host types selected from the possible values (Camel, Human). The variable *d*_*s*_ represents the duration in years between the first and last samples in the dataset. (Note that although priors are shared between types in this analysis, independent type-specific rates were estimated.)

| Parameter | Unit | Value/Prior distribution |
| --- | --- | --- |
| $R_{ii}$ | — | LogNorm(0, 0.5) |
| $R_{ij}$ ($i \neq j$) | — | Exp(1) |
| $\delta_i$ | Year<sup>-1</sup> | LogNorm(3, 0.5) |
| $s_i$ | — | Unif(0, 0.1) |
| $r_i$ | — | 1 |
| $T$ | Year | LogUnif($d_s$, 20) |

For the MCMC portion of the analysis, 5 independent chains of approximately 9 *×* 10^7^ iterations each were run, requiring approximately 90 hours of computation per chain. This was sufficiently long that all parameters of interest had an effective sample size (ESS) greater than 200. These chains were then compared to assess statistical convergence. (Supplemental Figure S4 shows close agreement between sampled distributions from independent chains, indicating convergence.) The first 10% of iterations from each chain were discarded to account for burn-in, and these truncated chains were then combined for further analysis. Subsequently, stochastic mapping of ancestral types and particle filtering of epidemic trajectories were conducted on one in every 10^6^ MCMC iterations, yielding 410 sampled edge-typed trees and multi-type trajectories in total. (Altogether, sampling these trajectories required approximately 30 minutes of additional computation, amounting to roughly 0.1% of the total analysis time.) Due to the large interval between MCMC iterations, we can regard these as effectively independent draws from the respective marginal tree and trajectory posteriors and thus sufficient to characterise these posteriors.

Figure 4 summarizes the results of this analysis. In Figure 4A, a maximum clade credibility (MCC) summary tree is shown with annotations indicating the ancestral host types best supported by the posterior edge-typed tree posterior. While minor differences exist, the same patterns observed by Dudas et al. (2018) using the structured coalescent model are also visible on this tree, with clear indication of a persistent camel-associated reservoir producing periodic spillovers into the human population.

**Fig. 4:**
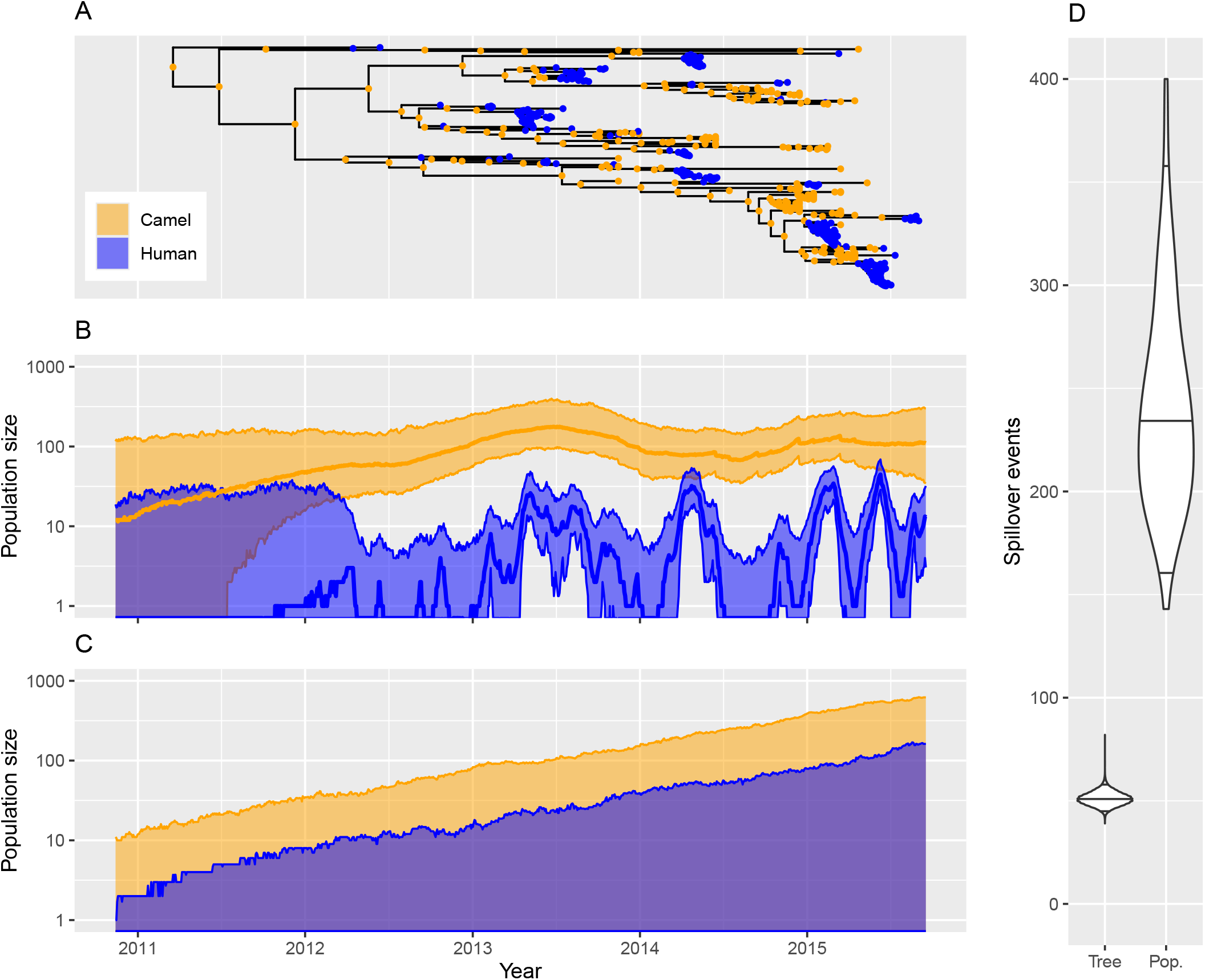
Joint inference of phylogenetic tree with ancestral host types and host-specific epidemic trajectories indicating the timing and frequency of MERS camel to human spillover events. (a) A maximum clade credibility (MCC) summary from the dated phylogeny posterior with most highly probable host types at internal nodes marked. (b) Marginal posterior of infected host population size dynamics sampled using our particle filtering approach, with the thick central lines representing the median and the bounds of the shaded areas representing the 2.5% and 97.5% quantiles of the marginal host-specific infected population size posterior at each time point. (c) Equivalent visualization of infected host trajectories simulated naively from the inferred model parameters, illustrating the importance of the tree in recovering infection dynamics. (d) Marginal posteriors of spillover events present on the tree (i.e. ancestral to included samples) and present in the larger population since 2011.

By counting events on the edge-typed trees, we found that the distribution of camel to human transmissions directly ancestral to the dataset has a 95% HPD of [45, 57] and median of 51 transmissions. These numbers are comparable to those determined by Dudas et al. (2018) applying the same approach to their structured coalescent results, who reported a 95% HPD of [48, 63] and a median of 56 for the number of transmissions.

The population-level spillover dynamics are shown in figure 4B, which displays a summary of the host-specific infected population size trajectory posteriors, including median values and 95% central posterior density intervals. The nearly constant case load in the camel population is clearly visible. (Interestingly, this displays strong similarities to the Bayesian Skyline Plot estimate of the camel-associated effective population size shown figure 5 of Dudas et al. (2018).) Additionally, the spillover outbreaks in the human population are reflected in the well-resolved spikes in the human-associated population trajectory.

For comparison, figure 4C summarizes a distribution of trajectories produced by directly simulating the birth-death process using model parameters sampled from the posterior. Unlike the trajectory distribution shown in figure 4B, this alternative distribution does not represent the true posterior distribution of population size trajectories, as it neglects the specific temporal information carried by the divergence time distribution of the phylogenetic tree, which in figure 4B is directly incorporated by our particle filtering approach. This panel makes clear the importance of taking this information into account, with this distribution displaying little ability to resolve any of the spillover features present in the true posterior distribution.

Importantly, the sampled trajectories can be regarded as sequences of model events. We used these to characterise the marginal probability distributions for the number of camel to human transmission events in the whole population from which our sample was drawn. The marginal posterior for the number of these events has a 95% highest posterior distribution (HPD) interval of [143, 421] and a median of 236. This is significantly higher than the number of spillover events occurring on the edge-typed trees and thus ancestral to the sampled sequences. A comparison of the marginal posterior distribution for the camel to human transmission events in the edge-typed tree with the corresponding distribution for events in the wider population is shown in figure 4D.

## Discussion

Multi-type birth-death phylodynamics inference methods have traditionally focused on the inference of the number, type and timing of events, such as lineage divergences and type-change events, directly affecting lineages ancestral to sampled sequences or individuals. By enabling estimation of type-specific population size dynamics, the method presented in this paper extends this capability to allow direct inference of the number, type and timing of events in the larger population which are not necessarily directly ancestral to the sampled genetic data. The approach is applicable to structured (multi-type) birth-death models for which thus far inference of population size dynamics was not generally possible. This follows on from our previous work (Vaughan et al. 2019) which used a particle marginal Metropolis-Hastings (PMMH) approach of Andrieu et al. (2010) to jointly sample both trees and population trajectory posteriors under a variety of linear and nonlinear single-type birth-death models.

As demonstrated in the MERS-CoV case study, the ability to impute these latent multi-type population trajectory variables has clear applications to the study of large epidemics, where the specific details of the subset of the population ancestral to the sample set is often not of direct interest. Instead it is usually the larger encompassing population of infected hosts that we wish to probe. Beyond epidemiology, there are also natural applications in other fields which make use of birth-death phylodynamic models. In macroevolution, models such as fossilized birth-death (FBD) together with linear multi-type models—BiSSE (Maddison et al. 2007), MuSSE (FitzJohn 2012), HiSSE (Beaulieu and O’Meara 2016), etc.—are commonly used to infer trait-dependent speciation and extinction rates from trees inferred using genetic and morphological data. Our approach provides an efficient and self-consistent route to estimating and testing hypotheses relating to the corresponding trait-dependent species richness dynamics under these same models.

Even when the phylogenetic tree and parameters such as birth, death and migration rates are the sole focus, inferred trajectories may still be useful in gauging the appropriateness of phylodynamic models. After all, as discussed in the introduction to this manuscript, birth-death-sampling processes imply the existence of such trajectories even when traditional methods report only the marginal distributions of the phylogenetic trees and parameters. Nonsensical properties of inferred trajectories may reveal existing but previously-obscured deficiencies in the ability of the model to adequately describe the intended population dynamics and sampling processes. Such deficiencies could also be assessed quantitatively as part of a formal model adequacy test in which summary statistics of inferred trajectories were compared with statistics computed via posterior predictive simulation. (Possible summary statistics for such a test might include type-specific population size bounds or counts of specific birth-death trajectory events.)

We have shown that estimation of multi-type population trajectories can be performed with relatively minor increase in computational effort relative to that of the MCMC inference which must still be performed in the manner described previously (Kühnert et al. 2016; Scire et al. 2022). However it is important to emphasize that the computational demand of the MCMC inference portion is significant, and is the rate-limiting step of the inference. In our experience, multi-type birth-death MCMC inference tends to be practical only for trees of hundreds of tips and for models with relatively few types Scire et al. (2022), although techniques have been developed which will help raise these practical limits in future (Louca and Pennell 2019; Zhukova et al. 2023).

We note that the effect of the restriction of the model space to linear models—which cannot, for instance, directly account for density-dependent shifts in transmission rate—can be somewhat ameliorated through the application of time inhomogeneous rate variation, which our approach supports via arbitrary piecewise-constant variation in rate functions, as shown by Kühnert et al. (2014). However, MacPherson and Pennell (2024) recently demonstrated that such approximations cannot fully account for the effects of density dependence on population size dynamics.

To conclude, we have presented a Bayesian approach for the inference of complete multi-type birth-death population size trajectories from genetic sequences. Our application of the method to MERS-CoV demonstrates that this can be used to directly infer interesting properties of populations from which genetic samples are taken, such as the host-specific infected population dynamics and the times and numbers of zoonotic events (rather than merely the number of zoonotic events ancestral to the genetic sample). Importantly, we have shown that extending existing MCMC-based phylodynamic analyses to incorporate trajectory inference can be accomplished with relatively little additional computational overhead. For these reasons, we believe that this new approach will prove to be a useful and practical addition to the phylodynamics tool set.

## Acknowledgements

The authors gratefully acknowledge Nicola Mulberry who provided valuable feedback on an early version of the manuscript, together with three anonymous reviewers who provided many helpful suggestions. This project has received funding from the European Research Council (ERC) under the European Union’s Horizon 2020 research and innovation programme grant agreement no. 101001077.

## Data availability

All of the BEAST 2 input files and associated scripts required to reproduce the analyses and results presented in this paper, together with corresponding usage instructions, are available for download from GitHub (https://github.coM/tgvaughan/MultiTyPeTrajectoryAnalyses). The method itself is implemented as a new package, BDMM-Prime, which is distributed as free software under the terms of the GNU General Public License version 3. The source code for the software is available from its GitHub repository (https://github.coM/tgvaughan/BDMM-PriMe), and a complete online manual including a detailed tutorial can be found at the package website (https://tgvaughan.github.io/BDMM-PriMe).

