## Supplemental text and figures. for "Bayesian phylodynamic inference of multi-type population trajectories using genomic data"

### Supplementary Material for "Bayesian phylodynamic inference of multi-type population trajectories using genomic data"

#### 1 Internal node transition probabilities

In order to determine the probability of a type transition at an internal node during the stochastic mapping process, consider the following situation.

An internal node in a tree consists of a parent edge $e$ and two daughter edges, $e_l$ and $e_r$. The base of the parent edge (just before the node) has type $s$. We aim to compute the following probability:

$$P(V(e_l,t_e)=s_l, V(e_r,t_e)=s_r|V(e,t_e)=s,\mathcal{T},\eta)\equiv W^e_{s,s_l,s_r}. \tag{1}$$

Expanding $\mathcal{T}$ as the union of $\mathcal{T}_\downarrow(e,t_e)$ and $\mathcal{T}_\uparrow(e,t_e)$, we have

$$W^e_{s,s_l,s_r}=P(V(e_l,t_e)=s_l,V(e_r,t_e)=s_r|V(e,t_e)=s,\mathcal{T}_\downarrow(e,t_e),\mathcal{T}_\uparrow(e,t_e),\eta). \tag{2}$$

The linearity of the MTBD process and the resulting independence of subtrees allows us to drop the dependence on $\mathcal{T}_\uparrow(e,t_e)$, giving

$$W^e_{s,s_l,s_r}=P(V(e_l,t_e)=s_l,V(e_r,t_e)=s_r|V(e,t_e)=s,\mathcal{T}_\downarrow(e,t_e),\eta). \tag{3}$$

We now define unnormalized transition probabilities

$$\tilde{W}^e_{s,s_l,s_r}=P(V(e_l,t_e)=s_l,V(e_r,t_e)=s_r,\mathcal{T}_\downarrow(e,t_e)|V(e,t_e)=s,\eta) \tag{4}$$

such that $W^e_{s,s_l,s_r}=\tilde{W}^e_{s,s_l,s_r}/\sum_{s_l's_r'}\tilde{W}^e_{s,s_l's_r'}$. Considering that $\mathcal{T}_\downarrow(e,t_e)$ can be decomposed into a birth event $\text{Birth}_{s,j}$ producing an individual of unspecified type $j$ together with left and right subtrees $\mathcal{T}_\downarrow(e_l,t_e)$ and $\mathcal{T}_\downarrow(e_r,t_e)$, we can rewrite this as

$$\begin{aligned}\tilde{W}^e_{s,s_l,s_r}=&P(\mathcal{T}_\downarrow(e_l,t_e)|V(e_l,t_e)=s_l)P(\mathcal{T}_\downarrow(e_r,t_e)|V(e_r,t_e)=s_r)\\&\times\sum_j P(V(e_l,t_e)=s_l,V(e_r,t_e)=s_r,\text{Birth}_{s,j}|V(e,t_e)=s)\\=&g^{e_l}_{s_l}(t_e)g^{e_r}_{s_r}(t_e)\sum_j P(V(e_l,t_e)=s_l,V(e_r,t_e)=s_r,\text{Birth}_{s,j}|V(e,t_e)=s)\end{aligned}$$

This last term is simply the probability density of the event $\mathrm{Birth}_{s,j}$ occurring at time $t$, combined with the probability of the event producing child lineages such that the left child has type $s_l$ and the right has type $s_r$. This probability is non-zero only for specific combinations of $s_l$, $s_r$ and $j$:

$$
\begin{aligned}
P(V(e_l, t_e) = s_l, V(e_r, t_e) = s_r, \mathrm{Birth}_{s,j} | V(e, t_e) = s) \\
= \begin{cases} \frac{1}{2}\lambda_{s,s_l} & j = s_l, s_l \neq s, s_r = s \\ \frac{1}{2}\lambda_{s,s_r} & j = s_r, s_l = s, s_r \neq s \\ \lambda_{s,s} & j = s_l, s_l = s, s_r = s \\ 0 & \text{otherwise} \end{cases}
\end{aligned}
\tag{5}
$$

where the factors of 1/2 account for a given birth event being compatible with two distinct orientations. Substituting this probability into the previous expression for $\tilde{W}^e_{s,s_l,s_r}$ and evaluating the sum over $j$ yields the result presented in the manuscript:

$$
\tilde{W}^e_{s,s_l,s_r} = g^{e_l}_{s_l}(t_e) g^{e_r}_{s_r}(t_e) \begin{cases} \frac{1}{2}\lambda_{s,s_l} & s_l \neq s, s_r = s \\ \frac{1}{2}\lambda_{s,s_r} & s_l = s, s_r \neq s \\ \lambda_{s,s} & s_l = s, s_r = s \\ 0 & \text{otherwise.} \end{cases} \tag{6}
$$

#### 2 Supplementary figures

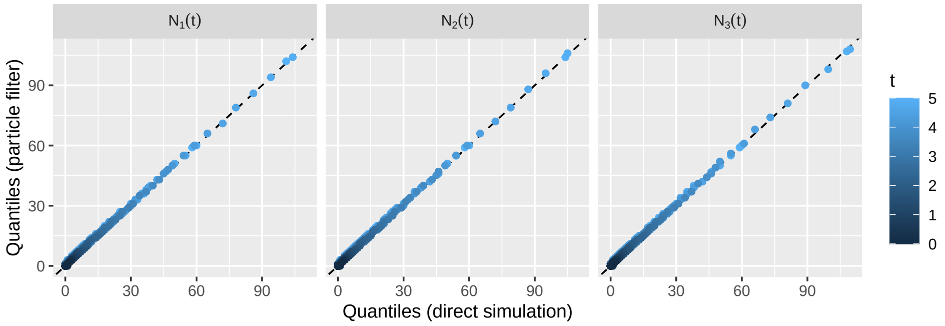


Figure S1: Comparisons between empirical quantiles of distributions of type-specific population sizes derived from trajectories generated by direct simulation and those generating using the particle filter, averaging over a broad family of three-type multi-type models as described in the manuscript. In contrast to the models considered in the manuscript, samples were generated by synchronous sampling events at times 4 and 5 following the start of the process at time 0. The type-specific sampling probability for each of these events was drawn from $\text{Unif}(0, 0.5)$. Each panel shows an overlayed series quantile-quantile plots comparing population size quantiles (10% through to 90%) for a type-specific subpopulation at a 51 evenly-spaced times spanning the simulation period. The dashed line is the diagonal indicating agreement between compared distributions.

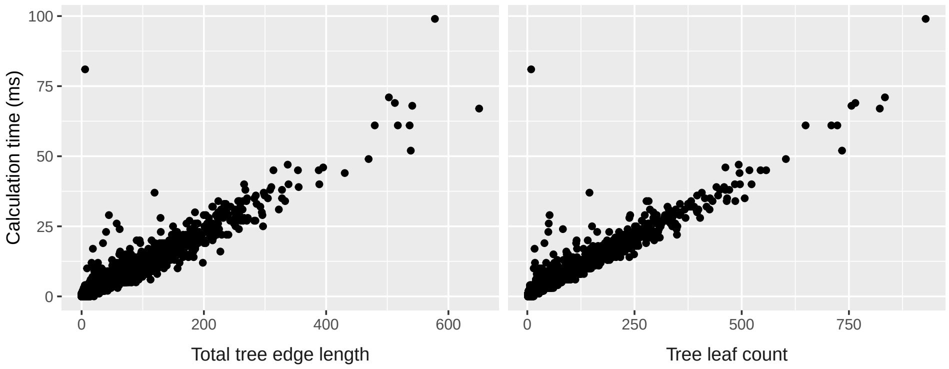


Figure S2: Empirical distribution of duration of stochastic mapping calculations performed as part of the stochastic mapping validation presented in the main text. The panel on the left shows these durations as a function of the number of leaves in the phylogenetic tree, while the panel on the right shows the same durations as a function of the total edge length of the tree. In both cases, the clear positive correlation highlights the importance of the expected number of generated type-change events in the time-complexity of the mapping algorithm.

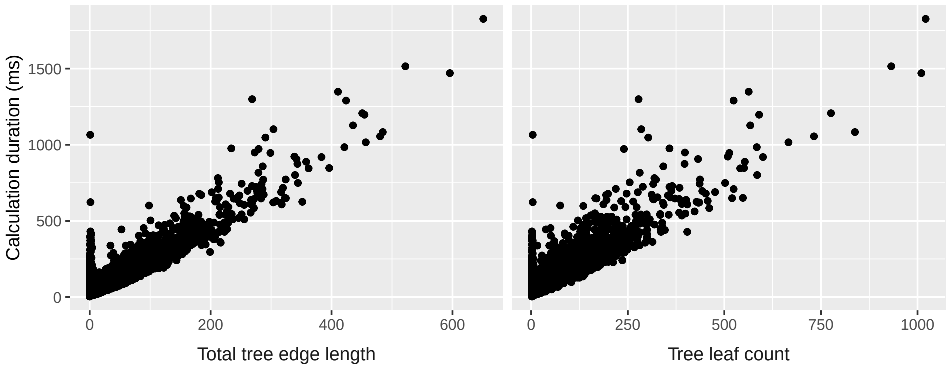


Figure S3: Empirical distribution of durations of trajectory inference calculations performed as part of the particle filtering validation presented in the main text. The panel on the left shows these durations as a function of the number of leaves in the corresponding phylogenetic tree, while the panel on the right shows the durations as a function of the total edge length of the tree. Both panels display an obvious positive correlation between the amount of time taken by the particle filter algorithm and the tree size.

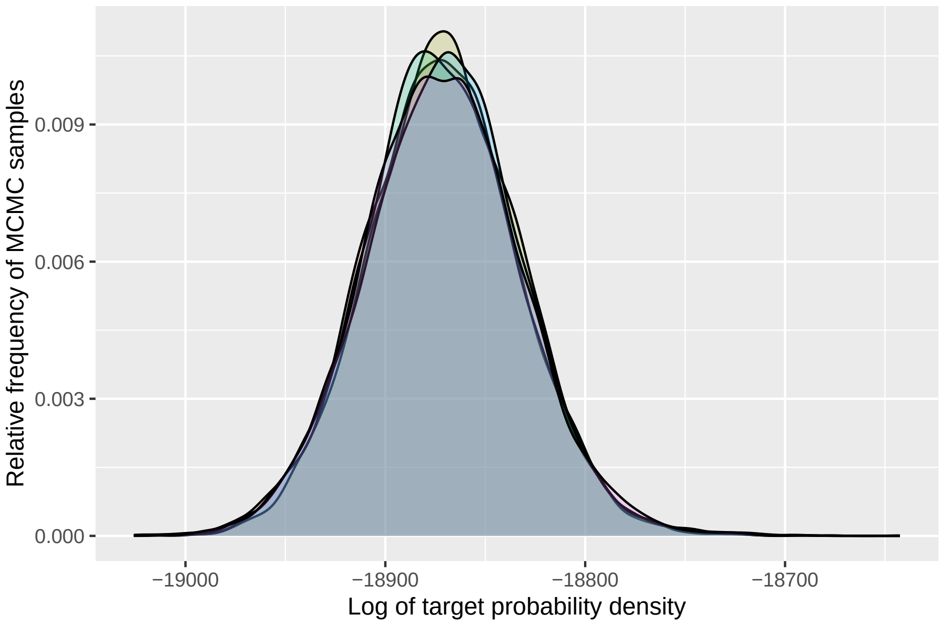


Figure S4: Comparison of relative frequency distributions of target densities generated by each of the 5 independent chains used in the MERS analysis. The agreement between these distributions provides evidence that the chains have equilibrated.
